# Current production as a rapid response expression reporter under micro-oxic and anoxic conditions

**DOI:** 10.1101/289140

**Authors:** Cody Madsen, Noelia Barvo, Ciara Fromwiller, Serenity Tyll, Brian Amburn, Danny Ducat, Bjoern Hamberger, Michaela A. TerAvest

**Author notes:** Author Contributions Conceptualization: Cody Madsen, Michaela TerAvest, Bjoern Hamberger, Brian Amburn, Noelia Barvo, Ciara Fromwiller, Serenity Tyll Methodology: Cody Madsen, Serenity Tyll Investigation: Cody Madsen Writing: Cody Madsen, Noelia Barvo, Ciara Fromwiller, Serenity Tyll, Brian Amburn Supervision: Michaela TerAvest, Danny Ducat, Bjoern Hamberger, Tim Whitehead.

## Abstract

Inducible gene expression is crucial for regulating cellular processes and production of compounds within cellular pathways. Yet, inducing gene expression is only the first step to utilizing cellular processes for an applied purpose such as biosensors. Detecting when gene expression occurs is central to understanding the overall mechanism of the process as well as maximizing the process. Fluorescent proteins have been established as the primary tool for detecting gene expression in inducible systems. This study proposes electricity production as an alternate tool in reporting gene expression. Using a model organism for electricity production, *Shewanella oneidensis* MR-1, current was shown to be an efficient reporter for gene expression and comparable to superfolder green fluorescent protein (GFP). Through regulation of the lac operator and T7 promoter, current production was induced by isopropyl β-D-1-thiogalactopyranoside (IPTG) addition. IPTG addition induced translation of GFP and the MtrB protein, which complemented a ∆*mtrB* strain of *S. oneidensis* MR-1 and restored current production. This inducible system generated reproducible current in 18 minutes in both micro-oxic and anoxic conditions. These results show that current is a fast reporter for gene expression.

**Financial Disclosure:** The team was supported by the following departments and colleges at Michigan State University: College of Natural Science, College of Engineering, Biochemistry and Molecular Biology Department and Plant Research Laboratory. The team also received support from the DOE Great Lakes Bioenergy Research Center (DOE Office of Science BER DE-FC02-07ER64494) and startup funding from the Department of Molecular Biology and Biochemistry, Michigan State University and support from Michigan State University AgBioResearch (MICL02454) (to B.H.). This work was also supported by NSF CAREER (Award #1254238) to T.A.W. MSU Alpha Chi Sigma also supported the team.

**Competing Interests:** The authors declare that no competing interests exist.

**Ethics Statement:** N/A

**Data Availability:** All data will be supplied upon request by the corresponding author.

This work was assessed during the iGEM/PLOS Realtime Peer Review Jamboree on 23^rd^ February 2018 and has been revised in response to the reviewers’ comments.

## Introduction

Past studies have shown that green fluorescent protein (GFP) is a useful tool for monitoring inducible gene expression since 1994. GFP can be fused to a gene of interest that cannot be measured as easily quantitatively. Placing GFP downstream from the gene of interest in an operon also generates accurate expression data regarding the gene of interest [1]. GFP has been used widely in synthetic biology as a reporter protein in various studies including monitoring phenol degradation, the production of anti-viral protein, and the production of interferons to prevent virus replication [2–4]. Given their results, fluorescent proteins have been a focus for gene reporter systems in many applications [5,6]. They are valuable molecular tools for *in vivo*, real-time, imaging or measurement of living specimens, but most are restricted to aerobic systems [7]. Mature fluorescent proteins can be seen and measured easily, making for a convenient experimental approach. Additionally, GFP has been utilized across multiple disciplines and even across several different organisms. Plants, microbes, yeast and even mammalian cells have been engineered to express GFP in order to indicate transcription of genes [6,8–10]. However, there are limitations of using GFP as a reporter protein because GFP requires oxygen for maturation and requires at least four hours for chromophore cyclization [11]. Therefore, GFP cannot be used as a reporter under anaerobic conditions. However, current measurements can be used as a reporter under anaerobic conditions [12,13]. GFP also may not be an ideal reporter for faster changes in promoter activity as GFP requires 4 hours at 22 degrees Celsius to form [11]. Yet, current as shown in this study shows sensitivity to induction within minutes. Current measurements show promise as an orthogonal reporter system for protein expression. Additionally, current can be measured in real time with a very high sampling rate to give nearly continuous data.

Recent studies have begun to investigate other means of reporting gene expression. One study has even shown the use of acoustic reporter genes, using gas vesicle production, which can be detected via ultrasound, as an output signal [14]. Accordingly, finding alternate tools for gene expression shows promise as a valuable avenue to researchers in multiple fields. One distinctive tool that has been developed in recent studies is using oxygen-independent, flavin-based fluorescent proteins as reporters under anaerobic conditions [7].

This study proposes that electricity production can act as another useful tool for reporting gene expression under anaerobic conditions. The marine bacterium, *Shewanella oneidensis* MR-1, was utilized as the model organism for electricity production. Yet, previous studies have shown that this electricity production can be transferred to other organisms such as *E. coli* [15]. The entire Mtr pathway was successfully transferred to *E. coli* and produced current [15]. Yet, studies are still being developed to optimize the transfer of the Mtr pathway to *E. coli* and eventually other organisms [16]. Therefore, the Mtr pathway has the potential to be utilized by other microbes in place of GFP as another system for reporting gene expression.

*S. oneidensis* MR-1 has been intensely studied for its ability to generate electric current via its metal reduction (Mtr) extracellular electron transport pathway [15,17]. The electron transfer pathway consists of four proteins: CymA, an inner membrane tetraheme cytochrome *c*; MtrA, a periplasmic decaheme cytochrome *c*; MtrC, an outer membrane decaheme cytochrome *c*; and MtrB, a transmembrane porin which stabilizes interaction between MtrA and MtrC [18] (Figure 1). These proteins are utilized during anaerobic respiration with solid metal oxides and electrodes [18–20]. Dehydrogenases oxidize electron donors such as lactate or NADH and transfer electrons to a membrane-soluble quinone. CymA accepts these electrons and transfer them to MtrA [13]. MtrA, a decaheme cytochrome, donates the electrons to the extracellular redox carrier MtrC in a process facilitated by the outer membrane porin, MtrB. Finally, MtrC can donate electrons to an external acceptor molecule, including the anode of an electrical circuit [13,17,18].

**Figure 1.**
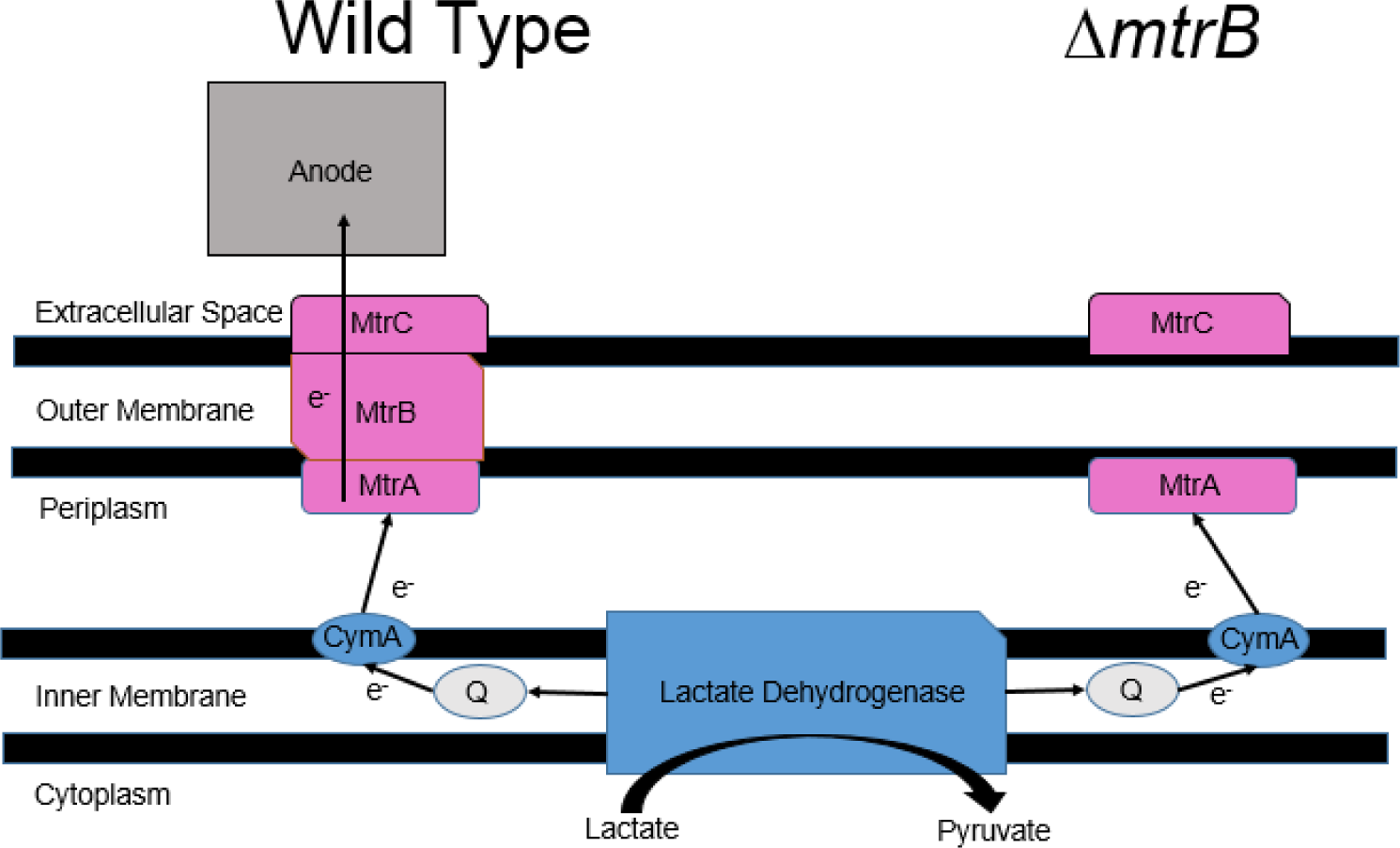
Modified Mtr pathway of *Shewanella oneidensis* MR-1. The wild type Mtr pathway allows for electron transport through MtrB to an acceptor. Electrons from lactate dehydrogenase are passed to a soluble quinone then to CymA which passes the electrons to MtrA. MtrA donates the electrons to MtrC facilitated by MtrB. MtrC then passes the electron to an acceptor. With the *mtrB* gene removed, a crucial gap forms in the native pathway stopping electron flow to the acceptor. After adding IPTG, MtrB is restored which results in increased current production measured at the anode.

**Figure 2.**
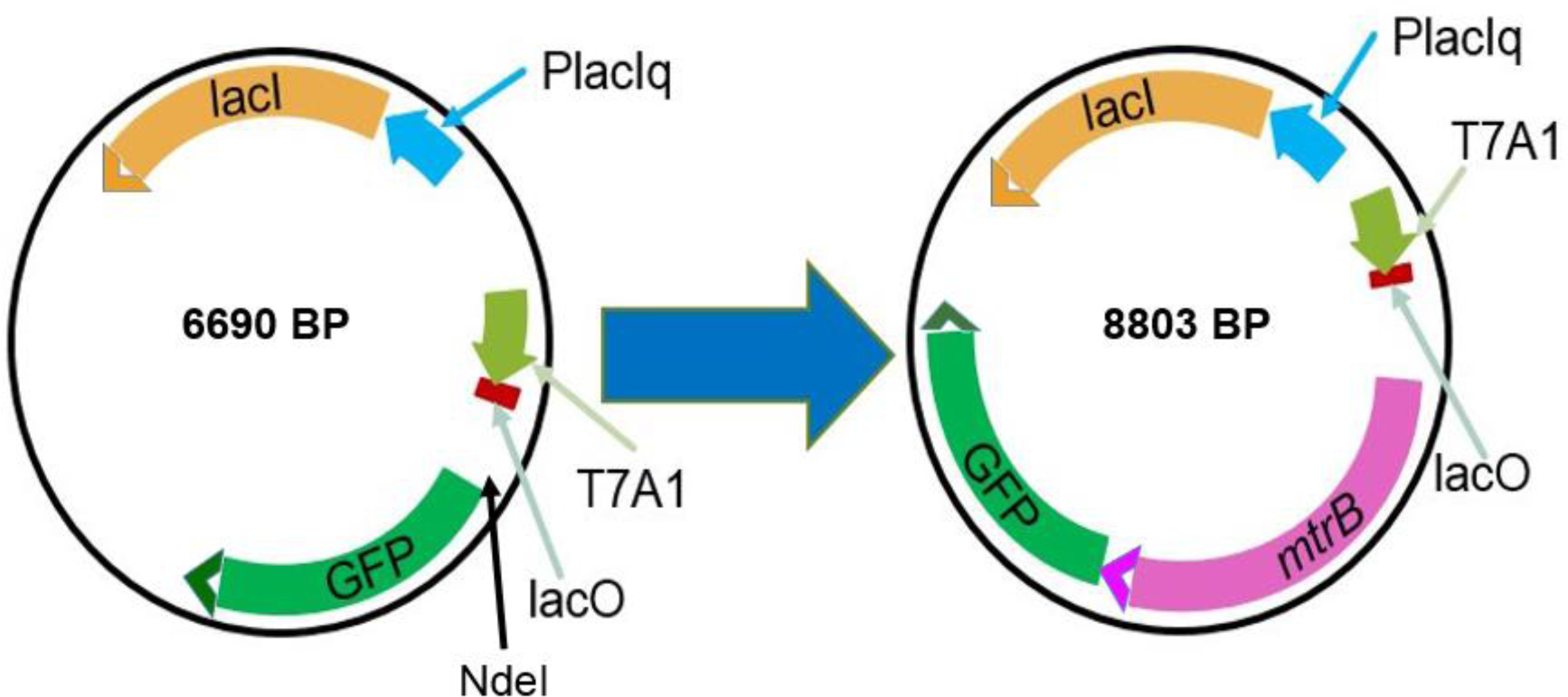
Plasmid construct. The *mtrB* gene was inserted into pRL814 (left) under regulation of the lac operon and the T7 promoter to make pRL814_mtrB.

Previous work has demonstrated that the Mtr pathway can be placed under inducible control by deletion of a component of the electron transport system, *mtrB*, and complementation of the pathway by introducing a plasmid with *mtrB* under control of an inducible promoter [12,13]. Accordingly, this inducible mechanism could potentially be used as a reporting system even under anaerobic conditions that are not amenable for the expression of GFP. Utilizing a ∆*mtrB* strain, the *mtrB* gene can be expressed under control of T7 or any other engineered promoter. In this study, we used the T7 promoter with a lac operator and repressor to control electricity production. Addition of isopropyl β-D-1-thiogalactopyranoside (IPTG), allows of the transcription of the *mtrB* gene which results in restoration of the complete native pathway leading to inducible current production in bioelectrochemical systems.

## Materials and Methods

### Bacterial strain, vector, and culture conditions

*E. coli* WM3064 was used for transformations and plasmids were conjugated from this strain into *Shewanella oneidensis* MR-1 Δ*mtrB* using an established protocol [21]. The Δ*mtrB* strain was obtained from Dr. Jeffrey Gralnick (University of Minnesota). The vector pRL814 was obtained from Dr. Robert Landick (University of Wisconsin-Madison) and was used for all gene expression. *E. coli* WM3064 was grown in LB medium (37°C, 275 rpm) with 30 µg/mL diaminopimelic acid (DAP) because this strain is a DAP auxotroph. *S. oneidensis* MR-1 Δ*mtrB* was grown in LB medium (30°C, 275 rpm, 12-16 hours) and strains carrying plasmids were grown in 50 µg/mL spectinomycin for selection.

### Plasmid Construction and Verification

The *mtrB* gene and its ribosome binding site (RBS) was amplified by PCR using the *S. oneidensis* MR-1 genome as a template. The *mtrB* gene was reintroduced by inserting it into pRL814 via NdeI digestion and ligation, resulting in an IPTG inducible expression system. The *mtrB* gene plus the RBS was inserted into the pRL814 plasmid downstream of the T7A1 promoter and upstream of GFP, leaving the RBS for GFP intact. The construct was designed to allow validation of protein expression through measurement of both current and GFP fluorescence. The plasmid was prepared for the *mtrB* insert via a NdeI digestion. Ligation with T4 DNA ligase was used to incorporate the *mtrB* DNA into the plasmid. The plasmid was transformed into *E. coli* WM3064 and then conjugated into the Δ*mtrB* strain. The constructs were verified by colony PCR and gel electrophoresis followed by Sanger sequencing.

### *Measurement of inducible* mtrB

M5 minimal media was prepared in 1L volumes by adding the following components to 880 mL of dH_2_O: 0.225 g potassium phosphate dibasic K_2_HPO_4_, 0.225 g potassium phosphate monobasic KH_2_PO_4_, 0.46 g sodium chloride NaCl, 0.225 g ammonium sulfate NH_4_SO_4_, 0.117 g magnesium sulfate heptahydrate MgSO_4_ * 7H_2_O, 23.8 g HEPES free ccid (23.8 g for anaerobic experiments), and 0.1 g casamino acids (Sigma-Aldrich). The pH was adjusted to 7.2 using 5 M NaOH (~ 7 mL). After sterilization in an autoclave, 1X Wolfe’s minerals (no aluminum) and 1X Wolfe’s vitamins, and 50 µg/mL of Spectinomycin antibiotic was added.

Bioreactors made out of Mason jars (250mL) allowed measurement of current output by consisting of a carbon felt electrode, a tungsten wire counter electrode, and an AgCl reference electrode made out of oxidized silver wire and KCl salt dissolved in heated agar. The carbon felt electrode was cut to 25 mm X 25 mm square then titanium wire was inserted through the felt and sealed with carbon cement. These electrodes were allowed to dry overnight in a fume hood. The counter electrode was assembled by inserting a titanium wire through the stopper then was surrounded by a plastic housing. The housing was used to contain and direct hydrogen gas production out of the bioreactor to prevent positive feedback by hydrogen oxidation at the anode. The plastic housing was created from a plastic syringe, punctured to allow ventilation. After adding M5 minimal media, all these elements plus a sampling needle were enclosed by a rubber stopper inside the Mason jar and the bioreactor was autoclaved. After autoclaving, the reference electrode was made by heating (~100° C) a combination of 50 mL dH_2_O and 0.75 g Bacto Agar in a 100 mL beaker, stirring at approximately 650 rpm. KCl was added until saturation (~23 g), then the hot solution was poured into a glass housing with a magnesia frit in contact with the reactor contents. A stainless steel needle was attached to a syringe, and inserted into the bottom of the housing to suction and remove any air to allow the solution to fill the cylinder in its entirety. An oxidized silver wire connected to a rubber stopper sealed the top of the housing to create the Ag/AgCl reference. The oxidized silver wire was prepared by applying a +0.2 V_Ag/AgCl_ to the silver wire while it was immersed in a KCl solution to deposit a layer of AgCl on the surface. The assembled Mason jars were placed on a low profile magnetic stirrer and the electrodes were connected to a lab-grade potentiostat (Bio-Logic Science Instruments, Part No: VMP3). Current production was measured using EC lab software (Bio-Logic Science Instruments, Version: 10.44). The software was programmed to the following settings to apply 0.200 V versus the Ag/AgCl reference[12]. M5 minimal medium was added to each reactor prior to sterilization in the autoclave. After allowing to cool, 1X Wolfe’s minerals, 1X Wolfe’s vitamins, 20 mM lactate (carbon source), and 5 µM riboflavin and 50 µg/mL spectinomycin were added to result in a final volume of 140 mL.

Baseline current was recorded for at least 2 hours prior to inoculation with *S. oneidensis* Δ*mtrB* containing the pRL814 plasmid (Δ*mtrB*_GFP) and Δ*mtrB* with pRL814 plus *mtrB* (Δ*mtrB*_GFP*mtrB*). These strains were run anoxically by bubbling nitrogen gas into the media inside the bioreactors. Then, Δ*mtrB*, Δ*mtrB_*GFP, and Δ*mtrB*_GFP*mtrB* were run under mostly oxic conditions as *S. oneidensis* MR-1 naturally converted the oxic conditions to slightly anoxic by biofilms. Bacteria were grown overnight, then diluted to OD_600_ equal to 0.01 in a final volume of 140 mL M5 minimal medium. After inoculation, current was monitored overnight, allowing bacterial growth background expression to equilibrate. After equilibration, *mtrB* expression was induced with 200 µM IPTG. This concentration was determined via to preliminary experiments. Subsequent current output was observed and recorded.

### GFP Maturation

Bioelectrochemical systems were sampled during current measurements for measuring GFP fluorescence. 1 mL samples were taken during the anaerobic and aerobic runs to allow for GFP to mature while inhibiting further production of protein. For the anaerobic samples, they were shaken at 500 rpm for an hour on ice under aerobic conditions. This allowed for the GFP to mature while slowing the rate of transcription/translation to minimize additional protein synthesis [22]. The samples were then transferred to a 96 well plate to be read on a plate reader. The aerobic samples were shaken at 500 rpm for one hour after addition of 10 µg/mL tetracycline. The tetracycline inhibits protein synthesis which allows for maturation of GFP and no further production of GFP as the healthier aerobic samples could synthesize more GFP even on ice [22,23].

### GFP Measurement

Cultures containing pRL814_mtrB were grown overnight in LB media + Spectinomycin antibiotic, then resuspended and diluted in M5 media to an OD_600_ of 0.01. Bacteria were then distributed in a 96 well plate in 200 µL volumes, such that three technical replicates for each strain (Δ*mtrB*_GFP*mtrB* Δ*mtrB_*GFP grown anaerobically and Δ*mtrB*, Δ*mtrB_*GFP, and Δ*mtrB*_GFP*mtrB* grown aerobically) and media background. A Bio-Tek, Synergy HTX Multi-Mode Reader took measurements for OD_600_ and GFP fluorescence simultaneously. GFP fluorescence was measured at 485 nm excitation and 527 nm emission. These measurements were taken sequentially using a clear-bottom, black 96-well plate.

## Results

### Impact of IPTG induction on current production

Figure 4 shows the difference in current between the background expression of the ∆*mtrB*_GFP*mtrB* strain and induction by IPTG. At 46 hours, 200 µM IPTG or an equal volume of sterile medium was added to each reactor. Current began to increase in the reactors that received IPTG ~18 minutes after injection, while current remained unchanged in the reactors that received medium (Figure 4B). The increase of current as shown by the blue line in Figure 4 shows induction of *mtrB* transcription by IPTG and also displays the ability of the cells to sustain current well after IPTG addition. These results are reflected in other experiments shown in Figures 5 and 6 by the characteristic increase in current initially then a decrease followed by induction via IPTG. Due to the *mtrB* gene being regulated by the lac repressor/operator on the pRL814 plasmid, background expression due to the equilibrium of the lac repressor occurred before IPTG addition.

**Figure 3.**
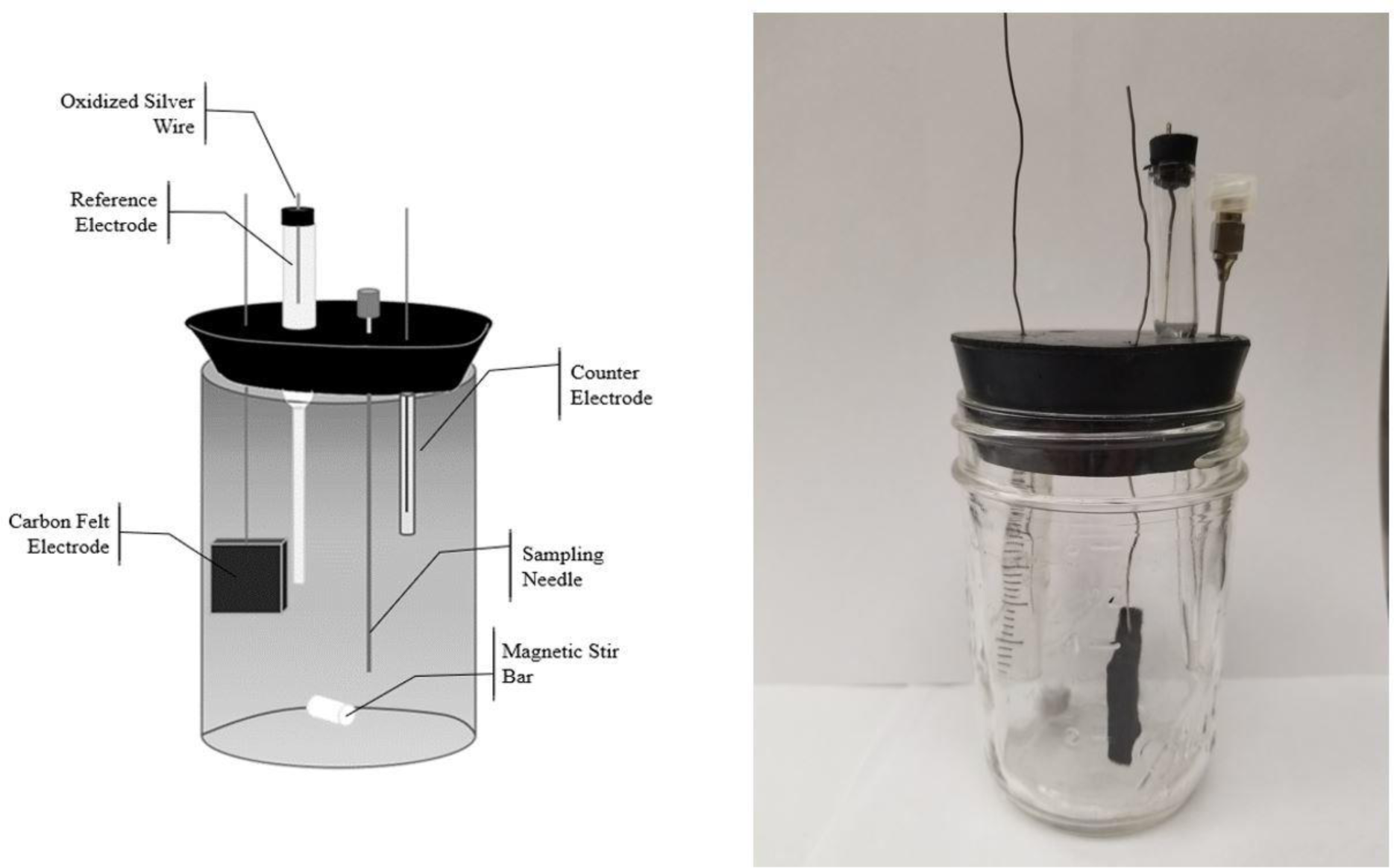
Engineered bioelectrochemical system. Completely assembled Mason jar bioelectrochemical system.

**Figure 4.**
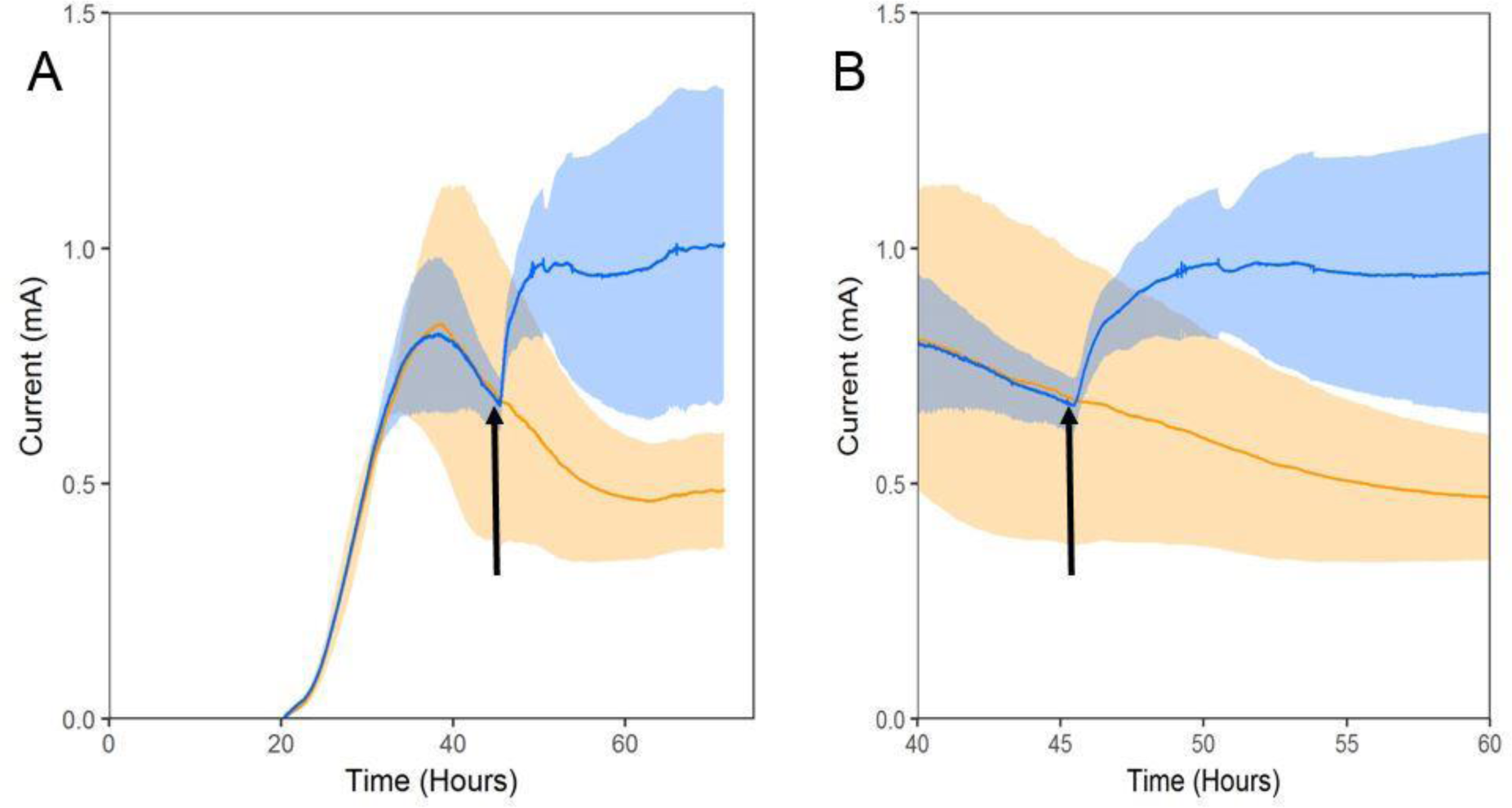
IPTG addition increases current production in modified strain. Current produced by ∆*mtrB*_GFP*mtrB* strain with addition of IPTG at time indicated by arrow (blue line) and ∆*mtrB*_GFP*mtrB* with addition of M5 media at time indicated by arrow (orange line) under anaerobic conditions. The shaded regions indicate standard deviations over multiple replicative runs. After 18 minutes post IPTG addition and media addition at 46 hours indicated by the arrow, current increases while the media addition makes no impact. Panel (B) shows a zoom in time scale at the point of IPTG induction.

**Figure 5.**
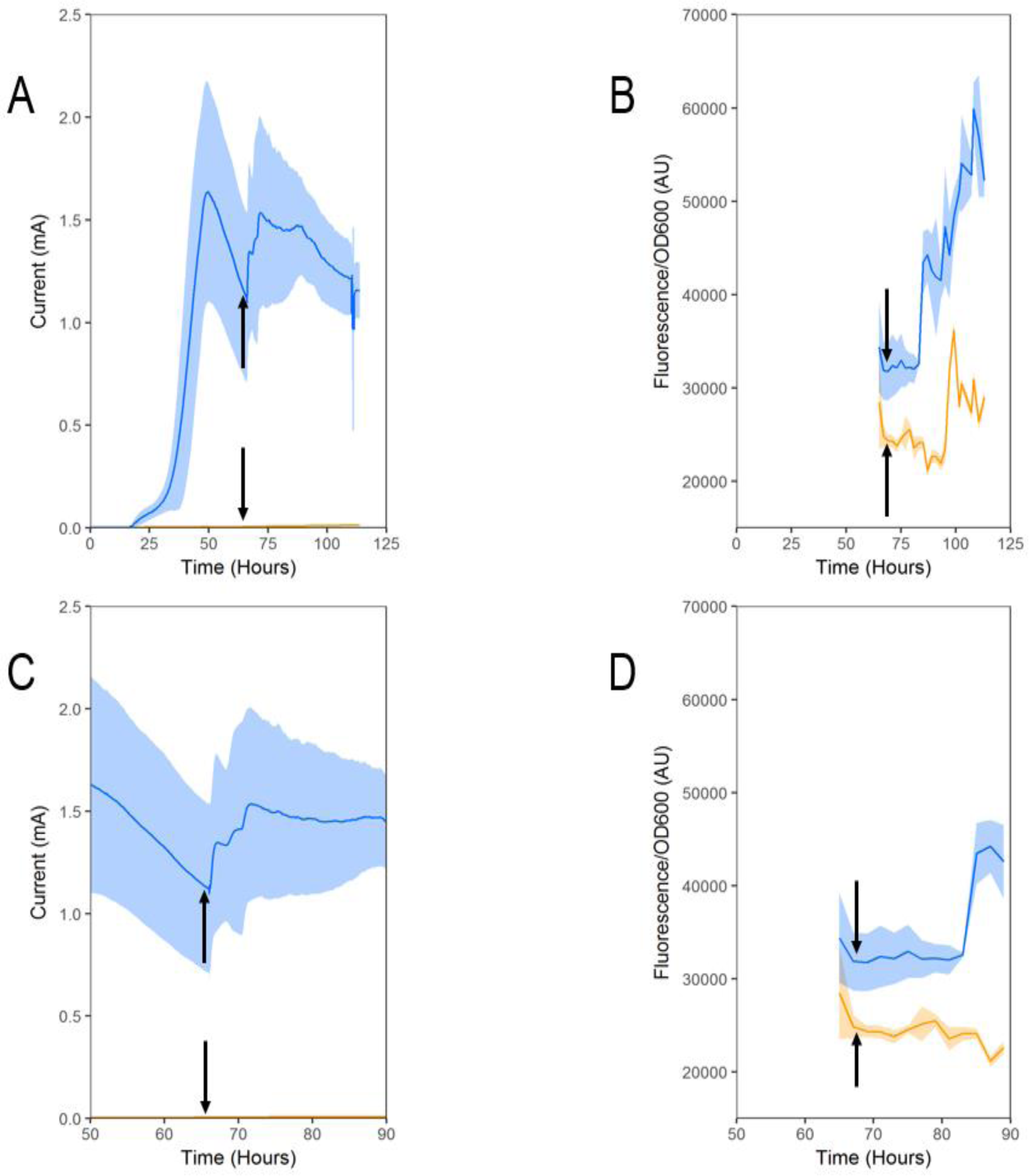
Anaerobic Current and GFP Fluorescence. Current production (A, C) and GFP fluorescence (B, D) produced by the ∆*mtrB*_GFP (orange lines) and ∆*mtrB*_GFP*mtrB* (blue lines) strains under anaerobic conditions. IPTG was added at 67 hours, as shown by all black arrows, to reactors for current and GFP measurement. Induction of the ∆*mtrB*_GFP*mtrB* (blue line) occurred 18 minutes later while the ∆*mtrB*_GFP (orange line) strain showed no induction (A, C). Induction of the ∆*mtrB*_GFP*mtrB* (blue line) occurred 17 hours later while the ∆*mtrB*_GFP (orange line) strain showed induction at 20 hours (B, D). Panels (C, D) display a zoomed in view on the induction time period.

**Figure 6.**
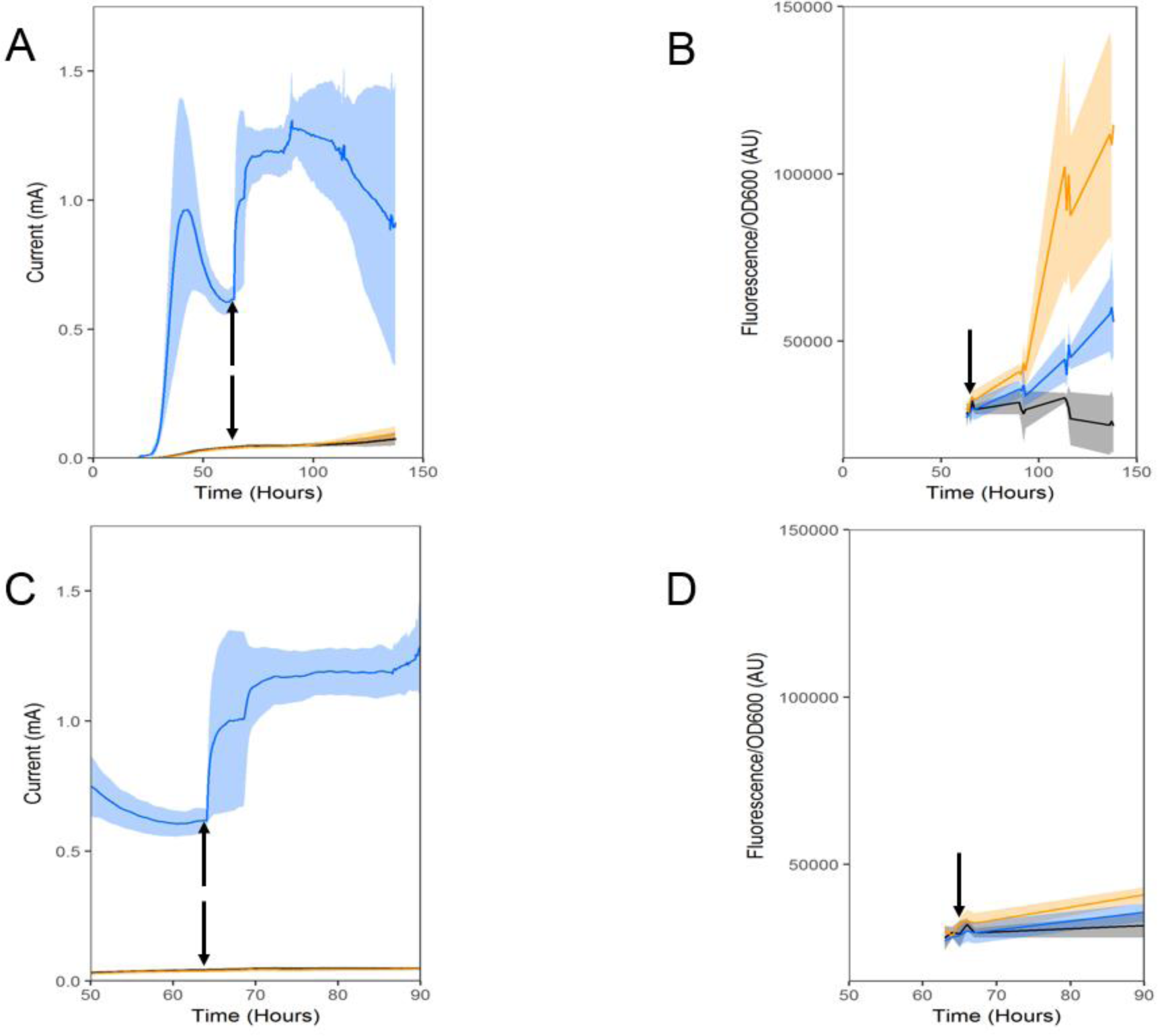
Aerobic Current and GFP Fluorescence. Current production (A, C) and GFP fluorescence (B, D) produced by the ∆*mtrB* (black lines), ∆*mtrB*_GFP (orange lines) and ∆*mtrB*_GFP*mtrB* (blue lines) strains under aerobic conditions. IPTG was added at 64 hours, as shown by all black arrows, to reactors for current and GFP measurement. Increase in current occurred 18 min later from the ∆*mtrB*_GFP*mtrB* strain while the ∆*mtrB*_GFP strain showed no induction (A, C). Additionally, ∆*mtrB*_GFP*mtrB* and ∆*mtrB*_GFP fluorescence showed significant induction at 90 hours (B, D). Panels (C, D) display a zoomed in view of the induction time period.

### Anaerobic Current and GFP Measurements

The ∆*mtrB*_GFP and the ∆*mtrB*_GFP*mtrB* strain were grown under anaerobic conditions and current was measured in triplicate (Figure 5A, C). Post inoculation at 18 hours, ∆*mtrB*_GFP began producing current with a maximum equal to ~0.02 mA. Conversely, ∆*mtrB*_GFP*mtrB* began producing background current that would equal approximately 1.5 mA. This background current reached a maximum at 50 hours, then began to decrease. At 67 hours the strains were induced with 200 µM IPTG. Following induction, ∆*mtrB*_GFP showed no increase in current while ∆*mtrB*_GFP*mtrB* began increasing in current 18 min after IPTG was added. The ∆*mtrB*_GFP*mtrB* continued to increase in current and reached a maximum 10 hours post IPTG addition.

Concurrently with current measurement, the bioreactors were sampled to measure GFP fluorescence of the two strains standardized against OD_600_. Following an hour of shaking on ice under aerobic conditions to mature GFP, fluorescence was measured (Figure 5B, D). Both the current and GFP fluorescence in Figure 5 follow the same time course, so IPTG addition occurred at 67 hours. Initial measurements were taken before induction to observe the background fluorescence level. GFP fluorescence increased minimally in the ∆*mtrB*_GFP*mtrB* strain (blue line) over the first several hours. The ∆*mtrB*_GFP*mtrB* strain then began to fluoresce significantly 17 hours following induction and fluorescence continued to increase until the end of the experiment. The ∆*mtrB*_GFP strain (orange line) began to increase 20 hours following induction but its fluorescence always remained lower than the ∆*mtrB*_GFP*mtrB* strain.

### Aerobic current and GFP measurements

Figure 6 shows the current production (A, C) and GFP fluorescence (B, D) of the ∆*mtrB*, ∆*mtrB*_GFP and ∆*mtrB*_GFP*mtrB* strains under aerobic conditions. The ∆*mtrB* (black line) and ∆*mtrB*_GFP (orange line) strains displayed residual current up to 0.1 mA while the ∆*mtrB*_GFP*mtrB* strain (blue line) showed higher background current production, then induction 18 minutes after IPTG addition at 68 hours. The background expression approximated at 1.0 mA around 40 hours based on Figures 4 and 5. Accordingly, once the current decreased all strains were induced with 200 µM IPTG and the ∆*mtrB*_GFP*mtrB* strain showed increase in current up to nearly 1.4 mA.

Just as in Figure 5 for the anaerobic conditions, the 1 mL samples were allowed to mature under aerobic conditions then these samples were measured for GFP fluorescence (Figure 6B, D). The GFP fluorescence was normalized to OD_600_ and plotted on the time course they were sampled which matches the time course of current production (Figure 6A, C). All strains were induced with 200 µM IPTG, as shows by all black arrows, and the ∆*mtrB*_GFP*mtrB* (blue line) as well as ∆*mtrB*_GFP (orange line) strains increased in GFP fluorescence. Minimal induction occurred over the first several hours. Approximately 26 hours after IPTG addition, both strains increased significantly with the ∆*mtrB*_GFP strain reaching a much higher maximum than the ∆*mtrB*_GFP*mtrB* strain. This is contrary to the anaerobic data which showed the ∆*mtrB*_GFP*mtrB* reaching a higher maximum than the ∆*mtrB*_GFP strain.

## Discussion

Inducible current production via IPTG due to the lac operon controlling transcription of the *mtrB* gene showed reproducible background expression and induction 18 minutes after IPTG addition. This is well within the timeframe of transcription and translation for a gene of this size [24]. Due to the strength of the T7 promoter and the equilibrium of the repressor, Figures 4, 5 and 6 display this consistent background expression. This background expression could be moderated by increasing expression of the Lac repressor or using a weaker promoter than the T7 promoter. However, as displayed in Figure 4, there is a clear distinction between background expression (orange line) and induction by IPTG (blue line). This result is repeated in Figures 5 and 6 under both anaerobic and aerobic conditions. Background current production follows the same trend for both conditions and the resulting induction shows consistency and reproducibility.

In our experiments, anaerobic current reached a higher overall level due to the electrode being the sole terminal electron acceptor, rather than competing with oxygen. Aerobic current was impacted by oxygen because it can act as a terminal electron acceptor while also improving the overall viability of the cells. Oxygen improving the viability of the cells could explain the sustainability of current over a longer period of time in Figure 6. Yet, *S. oneidensis* MR-1 began to develop characteristic biofilms at the interface between the media and the headspace under oxic conditions as early as one day into the run. This result indicates that the biofilms allowed for anoxic conditions within the media which promoted electron transfer to the electrode while the aerobic headspace supported viability of the cells. Again, these results begin to explain the sustainability of the current in Figure 6.

The current produced by the ∆*mtrB*_GFP strain under both anaerobic and aerobic conditions as well as the ∆*mtrB* strain under aerobic conditions demonstrates the impact of the Mtr pathway. These controls also show the effectiveness of manipulating this pathway to serve as a regulated reporting system. GFP fluorescence under anaerobic and aerobic conditions show the difference in response time to induction compared to current. While electricity responds in 18 minutes, GFP fluorescence under both conditions shows significant response only after several hours. This result may be explained by translation of the MtrB protein promoting a positive feedback for the cells by unlocking their native respiratory pathway while GFP production only places an added burden on the cells. This is further explained by supplementary Figures 1 and 2. The OD_600_ show that before induction the ∆*mtrB*_GFP*mtrB* strain shows substantial growth under aerobic and anaerobic conditions possibly due to background expression of the *mtrB* gene on the plasmid. Yet, after the induction, the cells gain back some fitness as their native pathway is unlocked which allows for the positive feedback to heighten sensitivity to the stimulus thus showing a rapid response to protein expression. Even though producing the MtrB protein also requires energy, the GFP protein production only burdens the cell without providing a benefit to the cell. This concept between the positive effect of MtrB versus the added burden effect of GFP could explain the sensitivity to induction displayed in Figures 4–6. Future experiments would look to quantify the sensitivity difference between MtrB induction and GFP induction through dose-dependence experiments. Yet, the sensitivity of current to induction provides an added benefit to this orthogonal, rapid response reporter for protein expression.

Reporters for gene expression are utilized in a variety of different disciplines. GFP has become the standard tool for this procedure due to its versatility in a variety of different organisms and applications. As seen in the results, this study proposes that electricity production can overcome some challenges in using GFP as a reporter for gene expression. Inducible current production shows reproducibility by repeatedly increasing 18 minutes post IPTG addition. This result occurs under both anaerobic and aerobic conditions, which overcomes the GFP deficiency of being restricted to aerobic conditions. Secondly, the rapid response of the induction improves on GFP response time in our system by over ten-fold. These two results can allow researchers to test for gene expression under anaerobic conditions in real time. Adding to the rapid response time, current can be measured easily by a potentiostat or Arduino board and the data can be viewed on a software program such as EC-Lab. This allows for the real time visualization of the induction and resulting impact on the cells. Additionally, this approach only requires a single chambered bioelectrochemical system that can be engineered from any sealable container that can be sterilized. This bioelectrochemical system attached to a simple potentiostat or Arduino board makes for an accessible way of measuring induction and therefore gene expression [25]. With GFP measurement techniques, constant sampling and measurement cycles are required to obtain the necessary data to know when gene expression is occurring. Our fast readout is beneficial when transcription of a gene can produce a compound toxic to the organism or when a useful compound such as a pharmaceutical or high-valued chemical is being synthesized by the organism [26]. The current would indicate when this compound is beginning to be produced and could be further engineered to indicate the amount of the compound being produced by the change in current.

Presently, this inducible system is functional in its native organism *S. oneidensis* MR-1 but could also be transferred to other organisms. The Mtr pathway has been successfully transferred to *E. coli*, which shows the potential of using this pathway outside its native host [15]. If the pathway was fully uncoupled from native metabolism, the response from inducing this system would be impacted solely by the inducer, resulting in an optimal reporter system. Yet, *S. oneidensis* provides a fast current read-out results from rapid oxidation of lactate. In this study, inducible current results were shown to be efficient at detecting protein expression using a lab grade potentiostat and simple bioreactors.

GFP has shown its versatility in being transferred to a variety of different prokaryotes and even into eukaryotic cells, including mammalian cells [6,8–10]. This versatility has promoted GFP to being the standard tool for gene expression because it is nearly universal to use across all organisms. While electricity production is centered in *S. oneidensis* MR-1, the promise of transferring this ability to other microbes could be essential to utilizing this inducible system to its fullest. As a result, microbes used to manufacture chemicals and pharmaceuticals or used in research to better understand specific gene expression are the main targets to utilize inducible current for gene expression monitoring.

## Acknowledgements

The authors acknowledge all faculty that provided specific assistance including: Dr. Robert Landick (plasmid), Dr. Jeff Gralnick (∆*mtrB* strain) and Dr. John Urbance (GFP microscopy). Additionally, the authors would like to acknowledge the postdoctoral researchers and graduate students Dr. Jefferson Plegaria, Dr. Bryan Ferlez, Erik Young and Donna Liebelt for help with the project. Furthermore, the lab manager Nicholas Tefft, M.S. and postdoctoral researcher Dr. Magdalena Felzak were pivotal in helping the team organize and execute experiments. Lastly, authors would like to recognize the companies and other supporters that supplied the resources needed including: Cayman Chemical, New England BioLabs and Dot Scientific.

