## Supplementary Materials for "Current production as a rapid response expression reporter under micro-oxic and anoxic conditions"

Supplementary Figures

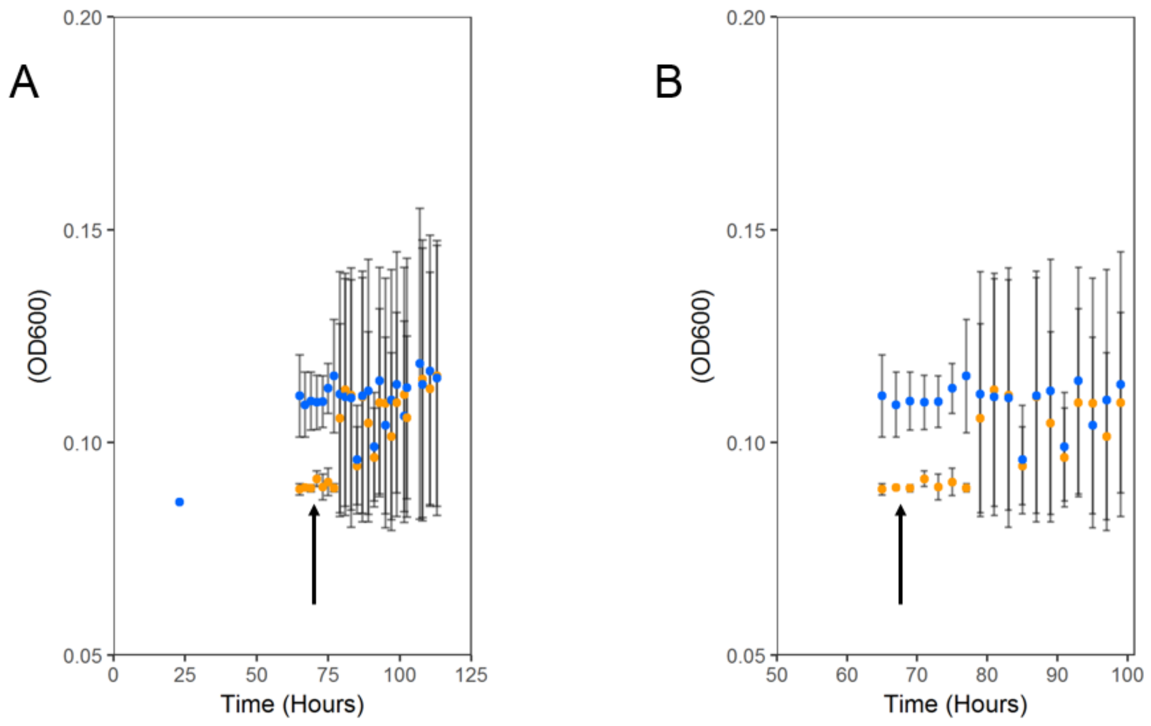


**Figure 1. Anaerobic $OD_{600}$**. $OD_{600}$ produced by the Δ*mtrB*_GFP (orange points) and Δ*mtrB*_GFP*mtrB* (blue points) strains under anaerobic conditions. IPTG was added at 67 hours, as shown by both black arrows, to reactors and $OD_{600}$ was measured alongside fluorescence. $OD_{600}$ shows little response to induction as cells appear viable before and after induction. Panel (B) display a zoomed in view on the induction time period.

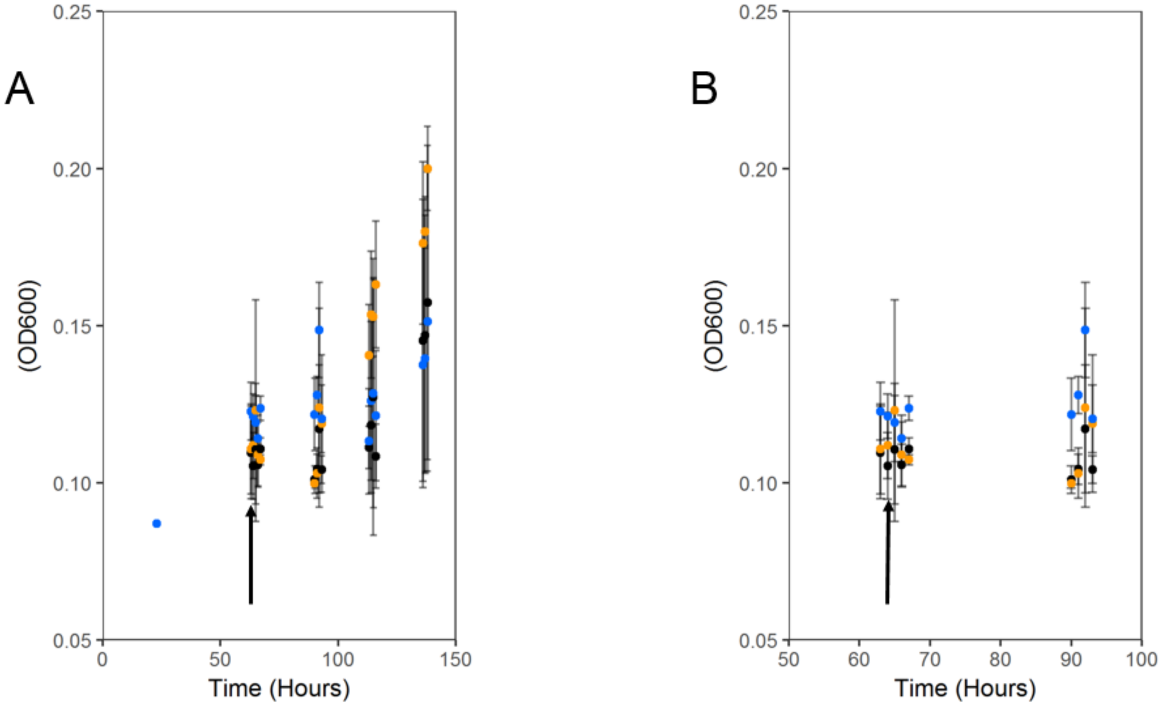


**Figure 2. Aerobic $OD_{600}$.** $OD_{600}$ produced by the Δ*mtrB* (black points), Δ*mtrB*_GFP (orange points) and Δ*mtrB*_GFP*mtrB* (blue points) strains under aerobic conditions. IPTG was added at 64 hours, as shown by both black arrows, to reactors and $OD_{600}$ was measured alongside fluorescence. $OD_{600}$ increases over time but shows no significant induction within the Δ*mtrB*_GFP*mtrB* strain. Background expression appears to give viability to the cells. Panel (B) display a zoomed in view of the induction time period.
