## Supplementary Materials for "Current production as a rapid response expression reporter under micro-oxic and anoxic conditions"

**Hi Cody, This reporter system appears to provide impressive reduction in response time - looks like it'll be a useful tool for the field. While I'm not going to focus on the results per se, here are a few suggestions for improving the message: - The title and abstract are not clear about what are you going to show us: especially regarding use under anaerobic vs aerobic conditions. Eg it's not clear whether you show that detecting current as proxy for expression can only occur under anaerobic conditions (where GF doesn't work so well) or can occur in both aerobic and anaerobic conditions (shown in figs 4-6?) The discussion is much more clearly written yet it is the title and abstract that will be focussed on by potential readers and need to attract readers to the key findings. - Use 'Rapid response' phrase in place of 'faster' where possible in title and abstract. - Perhaps this for a title edit: Current production via the Mtr pathway outperforms GFP as an expression reporter … I'll include other comments in-line.**

- We updated our title to: Current production as a rapid response expression reporter under micro-oxic and anoxic conditions
- We also updated the abstract to include micro-oxic and anoxic conditions as well

**Could you provide a link to a spread sheet containing with the data underlying your graphs in figs 4-6? Either deposit these in a repository or upload as supplementary files & cite these files in the results section - that way readers/reviewers can check and understand how you reached your conclusions.**

- We moved all files relating to each figure into file folders to be uploaded along with the submission. These include files for Fluorescence, current and $OD_{600}$

**I understand and appreciate that GFP is not suitable for anaerobic environments. My current project also has this constraint. However, fairly cheap UV lights can be used to show "glowing" colonies or streaks that are at least qualitative. Fairly simple and cheap fluorescent plate readers are more quantitative. And flow cytometry with fluorescent reporters enables a lot of measurements, and FACS enables sorting. I would focus this section on the anaerobic limitation of GFP, as well as the utility of having an orthogonal and totally different type of reporter - electric current generation, that could be used together with existing reporter systems, even in the same cells.**

- We agree with this comment that there are cheap devices available for GFP measurement so we removed this argument and focused on the prospect of using current under anaerobic conditions. We also wanted to suggest that current is a useful and orthogonal reporter for protein expression.

**I suppose to use this reporter in an organism that does not normally generate current, you'd have to install the whole Mtr pathway, with one gene like mtrB under control of the promoter of interest, vs. just expressing a single protein in the case of fluorescent proteins. So that's a disadvantage, but still it could be worth it for many researchers.**

- We agree that transferring the entire Mtr pathway is necessary and difficult. Yet, we added in more citations in the intro and discussion showing examples of how this has been accomplished already.

**You want to say that the mtrB gene can be inserted after any promoter of interest, to evaluate that promoter and the conditions under which it is active for gene expression. You demonstrate this with a specific example of lac and IPTG. But many others would use it differently and more generally.**

- We changed our discussion in this section to specifically discuss how we used the T7 promoter and a lac operator in our study to indicate that other studies could use any other promoters "...the mtrB gene can be expressed under control of T7 or any other engineered promoter. In this study, we used T7 promoter with a lac operator and repressor"

**The final sentence here is missing something? Important to get this introductory schematic correct!**
**Your last sentence needs to say that restoration of mtrB by complementation, under control of the promoter of interest, restores current production, which is measurable at an anode surface.**

- We updated our final sentence in Figure 1 to clarify the transfer of electrons to the anode via current restoration

**Would be good to show ORF directions of GFP and mtrB with arrows. Is it a little operon of the two? What kind of RBS site does each gene have?**

- We updated our Figure 2 to include new arrows that are different than the arrows on the promoters to distinguish the genes from promoters. We also added in our methods section where the RBS sites are located

**Bio reactors made out of mason jars – cool! Why don't you upload this technique and figure 3 on protocols.io. Once uploaded you can provide a link to it within this methods section - and use the link whenever you need to reference the technique going forward. You'll also get credit every time someone else cites the link. If you haven't already then check it out the possibilities for all your protocols here https://www.protocols.io/**

- Thank you for the offer! We considered the offer and decided we would like to keep this paper as the repository for the method in assembling the bioreactors.

**It's really interesting seeing the photograph alongside the labelled figure and should be a good aid to reproducing this study.**
**Agreed - nice figure. Critical to understanding the paper and very helpful too.**

- Thank you for the generous comments! We are glad that Figure 3 gives clarity to the project!

**hmsrocinante2112: Mention that panel B is a zoom-in of the same data in panel A. I assume the shaded regions represent something like confidence intervals or standard deviations. Please mention what specifically they represent (like error bars, you always need to define what specific statistic they represent).**

- We mentioned in all Figures that some panels are zoom-in on the induction time periods. We also mentioned that our shaded regions are our standard deviations after multiple replicate runs.

**This data is impressive, but given the message of the paper and the mention of induction after only 18 minutes, perhaps another graph can be added which 'zooms' in on the region of induction: for example, the scale of from 65 to 70 hours. This could further illustrate just how much quicker induction is when the reporter is current, not fluorescent.**

- We agree with this and added in zoom-ins in all of our figures for both current and fluorescence.

**Does low expression of mtrB make the cells very sick or slow to grow under anaerobic conditions? Is that a primary way they get energy under these conditions? I would be curious to know the OD600 vs. time trace for this. If cells are very sick when mtrB is not induced, that is not a great feature for a reporter system - when fluorescent proteins are not induced, the cells are arguably more fit, at least by a little bit. It would be good to give an honest assessment of this to the audience. Either lack of mrtB has little effect on cell health, which is good for a reporter system, but doesn't support your "energentics" argument below in trying to explain your GFP levels, or lack of mrtB has a strong negative effect on cell health, which is very bad for a reporter system, but might be consistent with your explanation of GFP levels.**

- We added two supplementary figures that show the $OD_{600}$ for both anaerobic and aerobic conditions. We also added to the end of the first discussion paragraph indicating that MtrB protein can serve as a positive feedback compared to a negative burden of GFP which allows for a heightened sensitivity to stimulus and a rapid response to protein expression

**I'm not convinced that you can explain the higher GFP levels in the ΔmtrB_GFP strain vs. the ΔmtrB_GFPmtrB strain by invoking "energy expenditure". Order of transcription in the operon, if that's the structure of the plasmid, could be a better explanation (there are many refrences for this, generally, the 3' end of mRNA gets degraded first). Plus, the levels of GFP are reversed in Fig 5 (anaerobic) vs. Fig 6 (aerobic), as you noted. After this point, there is a lot of conjecture in this Discussion that is not substantiated by your results, so you would need to tone down the speculation or do many more experiments or cite very relevant references. In the cells with the full Mtr pathway, under anaerobic conditions, does that pathway substantially increase the energy available to the cells? Can you show that?**

- We agree with this review that there is too much conjecture and have removed this point from the discussion.

**I follow your argument about the possible impact that mtrB can have anaerobically vs. its limited impact aerobically, but I and many readers don't know enough about how this microbe gets energy to know if it makes sense. Can you cite some other research on mtrB deletions and their effect on anaerobic growth of S. oneidensis? If you could change the order of GFP and mtrB in your operon, that would let you test whether that has a big impact or not, regarding your "energy" hypothesis related to transcription.**

- Again, we agree with this continued point and we have chosen to remove it from the discussion.

**Hi Michigan State University, Thanks for your submission! You have shown that your current based system is faster at detecting gene expression compared to GFP, can you compare the sensitivity of the two?**

- We mentioned in the discussion at the end of the first paragraph that in future we would want to compare the sensitivity by dose-dependent experiments

**Impressive work, very interesting idea. I am curious if you have tried to model the expression of the mtrB gene - is 18 minute induction what you would expect, given that some delay is necessary to transcribe, translate, and translocate the protein? Regardless, it is nice to see a reporter system with a response time that is actually within the generation time for a bacteria!**

- We have added a reference to the BioNumbers database indicating that the induction we observed is feasible.

**Thanks for this paper! I enjoyed reading it and learned several things. To summarize my most important comments from above, I think the readers need to see what effect the mtrB deletion has on cell growth and viability under anaerobic conditions. You most likely have the OD600 curves for induced vs. uninduced, so you should show that in a figure. If the uninduced cells have poor growth and viability, as I suspect they might, that is not a good feature for a gene expression reporter system. However, there is still hope that this system could be useful in general. If the whole Mtr pathway can be ported to an organism that does not need it or depend on it for viability, under whatever conditions (anaerobic or aerobic), then it could be used to monitor electrical current as a measurement for gene (mtrB) expression by a promoter of interest. That feature - neutrality with respect to cell viability, is a pretty important feature for the adoption and use of a reporter system. If the system described here is not neutral in this organism (S. oneidensis), it might be when ported to another, like E. coli or yeast or Clostridium, etc. I would like to see this type of material covered in the Discussion. Thanks again for sharing this work!**

- We have included the supplementary figures that show that the cells show fair viability before induction and gain more viability as a sensitive positive feedback. We also mentioned in the later parts of the discussion the onset of moving this entire pathway to *E. coli* to uncouple this pathway from native respiration. This will allow for a “cleaner” vector for biosensor experiments even though the process used in this experiment shows promising sensitivity for protein expression detection.
